# Genome-wide association reveals novel insights into the molecular mechanisms regulating stem volume in *Pinus taeda*

**DOI:** 10.1101/2022.12.06.519371

**Authors:** Alexandre Hild Aono, Stephanie Karenina Bajay, Felipe Roberto Francisco, Anete Pereira de Souza

**Author notes:** **Corresponding Author:** Anete Pereira de Souza. These authors contributed equally to this work (Alexandre Hild Aono, Stephanie Karenina Bajay, Felipe Roberto Francisco).

## Abstract

*Pinus taeda* (loblolly pine [LP]) is a long-lived tree species and one of the most economically significant forest species. Among growth traits, volume is the most widely considered trait in tree improvement programs. However, deciphering the genetic variants responsible for growth trait variations in conifers, such as LP, is particularly challenging due to the vast size and intricate complexity of *Pinus* genomes. We present a comprehensive genetic analysis of LP, focusing on markers associated with stem volume variation, to elucidate the molecular mechanisms governing high-performance phenotypes. We used a population of 1,692 individuals phenotyped for stem volume and genotyped these individuals using sequence capture probes. To conduct genome-wide associations, we utilized both genome-wide association study (GWAS) analysis and machine learning (ML) approaches. The markers identified in association with volume were found to be linked with the genes assembled from three distinct transcriptomes. These genes were subsequently used to construct gene coexpression networks, and through topological evaluations, we identified key genes with potential regulatory roles within stem volume configurations. Using a set of 31,589 SNPs, we defined 7 GWAS-associated SNPs and 128 ML-associated markers, all of which were correlated with multiple genes involved in diverse biological functions. Gene coexpression analysis revealed a group of 270 genes potentially associated with the regulation of genetic material. Key genes directly implicated in the regulation of growth and response to stress were identified, and inferences about their impact on pine development were subsequently elucidated. Our study not only offers insights into SNPs associated with stem volume but also elucidates a subset of genes characterized by unique regulatory features. These findings significantly advance our understanding of the genetic factors influencing growth traits, reveal candidate genes for future functional studies, and contribute to a broader comprehension of the genetic architecture underlying volume traits in LP.

## 1. Introduction

Since the 1950s, forest product companies have intensively invested in tree breeding, plantation silviculture, and forest management to meet the global demand for wood and fiber [1]. *Pinus taeda* (loblolly pine, LP) stands out as one of the most economically significant forest trees worldwide [2,3], particularly within the timber, pulp, and paper industries in the southeastern United States (US). Conifers, such as LP and other commercially grown tree species, are still considered largely undomesticated despite more than six decades of breeding. There has been increasing pressure to improve this species to increase productivity, forest value, and ecosystem services [4,5].

A single breeding cycle of LP using traditional phenotypic selection requires a substantial time investment ranging from 15 to 30 years, including at least 8 years dedicated to field testing [6]. Although classical pine breeding methodologies have made substantial progress in recent decades, the incorporation of biotechnological and omics-based approaches is a promising avenue for accelerating and expanding genetic gains [7–10]. Despite the successes observed in other plant species, the application of these strategies in LP and other conifers faces challenges primarily attributed to the complex genomes of these species.

Even with recent advances in sequencing technologies, the assembly of the pine genome has been constrained to a highly fragmented draft reference, consisting of more than one million scaffolds and encompassing up to 82% of various highly repetitive DNA elements [11]. This limitation is attributed to the intricate, poorly characterized, and extremely large genome of the pine species (1C=21.6 Gb) [12–14]. The inherent complexity of the LP genome has prompted researchers to primarily focus on restricted genomic analyses, predominantly on nuclear gene sequences, which have served as the primary source of molecular data for enhancing the improvement efforts for this conifer [15–18]. In this context, the inclusion of additional molecular data sources can support the improvement of LP, especially for highly quantitative traits of commercial importance, such as stem volume (SV).

The targeting of genetic breeding programs to produce varieties with high SV is prioritized for most cultivated tree species, including LP [19,20]. However, the achieving increases in volume based on molecular data is hampered by the polygenic nature of this complex trait [21]. Statistical methods for associating genotypes with phenotypes have been of great interest for breeding, especially as tools for genomic selection (GS) and for deciphering causative loci via genome-wide association studies (GWASs). These models encompass a variety of methods with varying degrees of complexity, computational efficiency, and predictive accuracy [22–24].

GWAS models rely on the estimation of genetic effects using many molecular markers, with the goal of capturing the phenotypic contributions from both coding and noncoding regions of the genome [25]. However, this framework has been insufficient for elucidating the evolutionary forces responsible for maintaining the variation in the complex genomes of coniferous trees. Despite the integration of extensive phenotypic and genotypic datasets with robust statistical models, the genetic markers identified through GWAS often reside outside annotated gene boundaries and may be relatively distant from the actual causal polymorphisms [26].

To achieve a more comprehensive understanding, additional methodologies that enable the elucidation of unknown molecular interactions must be developed. Furthermore, the integration of diverse omics layers, encompassing genomics, transcriptomics, and proteomics, can contribute to a more holistic perspective, uncovering intricate relationships within the genome. This multidimensional approach is essential for revealing the complex forces that govern the wide-ranging genomic variations observed in coniferous trees.

In this study, we employed a targeted genotyping approach to investigate the molecular forces influencing SV variation in a cohort of 1,692 individuals from a breeding population of LPs. Our primary objective was to identify molecular markers associated with SV through the integration of GWAS and tree-based machine learning (ML) algorithms. This comprehensive analysis aimed to determine the genetic architecture that underlies the volume variation across seven distinct LP populations. Furthermore, the selected markers were mapped to specific genes, providing valuable insights into the functional implications of these polymorphisms. These results were subsequently integrated into a gene coexpression network, providing evidence of the impact of volume variation on plant growth.

## 2. Materials and Methods

### 2.1. Breeding Population and Plant Material

Phenotypic data were collected during the 2^nd^ cycle of breeding and selection conducted by the Cooperative Forest Genetics Research Program (CFGRP) at the University of Florida. This population was generated through crosses of twenty-eight LP progenitors, which were used as females and males. A total of 25,080 full-sib trees, originating from 308 families (∼81 individuals per family), underwent field testing at seven distinct sites established in 2006. The experimental design followed a randomized complete block design, with single-tree plots organized in an alpha lattice containing 20 incomplete blocks in 12 replications with 300 trees.

The stem height (SH) and diameter at breast height (DBH) at 1.4 m were measured at 3 and 4 years of age, converted into decimeters (dm), and used for estimating the SV in dm^3^ based on the inside-bark total stem equation [27]:

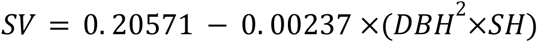

### 2.2. Phenotypic Data Analysis

The phenotypes were analyzed using ASReml v.2 [28] according to the following linear mixed-effects model for each environment:

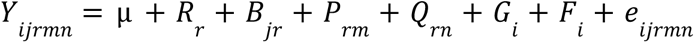

where *Y*_*ijrmn*_ is the SV measure of the *i*-th genotype within the *j*-th block, the *r*-th replication, and the *m*-th and *n*-th positions (rows and columns). The fixed overall mean is represented by µ, and the fixed effects of the *r*-th replication are represented by *R*_*r*_. The random effects were modeled to estimate the contributions of (i) the *j*-th block in the *r*-th replication (*B*_*jr*_); (ii) the *m*-th row in the *r*-th replication (*P*_*rm*_); (iii) the *n*-th column in the *r*-th replication (*Q*_*rn*_); (iv) the *i*-th genotype 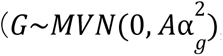, with 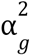 representing the genetic variance and *A* the pedigree-based relationship matrix); (v) the family of the *i*-th genotype (*F_i_*); and (vi) the residual error 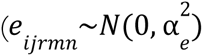, with 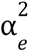 representing the residual variance). We excluded phenotypic measurements from trees that had died and calculated the average of the adjusted predicted least square means for each genotype.

All remaining analyses were conducted using R statistical software [29]. The individual narrow-sense heritability (*h*²) was calculated as the ratio of genetic variance 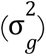 to the total phenotypic variance. The dominance ratio (*d*²) was determined by dividing the variance in the interaction between the block and replication by the total phenotypic variance. Term 4 was included to account for the presence of half-siblings within the block and for replication.

### 2.3. Genotypic Analyses

Genomic DNA was extracted from 1,692 LP seedling samples derived from 45 different crosses and grown at four distinct field sites. For genotyping, we employed sequence capture technology from Rapid Genomics (Gainesville, FL). From a preexisting collection of 54,773 probes, we selected 11,867 probes to represent 14,729 *P. taeda* unigenes [30]. The resulting target-enriched libraries were sequenced using the Illumina HiSeq 2000 platform.

The raw reads from the sequence capture data of the genotypes were aligned to the LP reference genome v.2.0 (GenBank accession GCA_000404065.3) [14] using Bowtie2 v.2.2.9 [31]. For read mapping, we considered only the regions surrounding the probes, extending 500 base pairs (bps) on each side of the sequence. SNP variant discovery was performed with FreeBayes v1.3.1 [32]. Subsequently, we employed VCFtools [33] and BCFtools v.1.3.1 [34] for SNP filtering. Our filtering criteria included selecting only biallelic SNPs and removing variants with a phred-scaled quality score less than 20, a minimum alignment score of 30, and allele frequencies ranging from 0.05 to 0.95. Additionally, we limited the proportion of individuals with missing information per SNP to a maximum of 10%. Any missing values for each SNP were subsequently replaced with the mean of the observed values.

Principal component analysis (PCA) was also conducted using the snpReady R package [35]. We estimated linkage disequilibrium (LD) between loci by calculating squared Pearson correlation coefficients (*R*²) for markers located within the same scaffold. LD decay was assessed using the exponential model *y* = *a* + *be*^(−*cx*)^ [36], where *x* and *y* represent the physical distance in base pairs and *R*², respectively. In this model, *a* + *b* represents the mean level of disequilibrium for loci at the same genomic position, and *e* represents the exponential term.

### 2.4. RNA-Seq Analysis

To assess transcripts within probes and potential annotations, we employed three independent experiments with samples from different pine species: (i) *P. taeda* (Pta) [18], (ii) *P. elliottii* (Pel, slash pine) [17], and (iii) *P. radiata* (Pra) [15].

In experiment (i), RNA sequencing (RNA-Seq) was performed on secondary xylem tissues from three 15-year-old trees, yielding three cDNA libraries obtained from the Yingde Research Institute of Forestry in Guangdong Province, China. The Illumina HiSeq 4000 platform was used for sequencing, as described by Mao et al. [18].

For experiment (ii), RNA-Seq was conducted on 16-year-old Pel trees, resulting in 16 cDNA libraries. These trees were sourced from Irani Celulose in Balneário Pinhal, Brazil. Sequencing was accomplished using the Illumina HiSeq High-Output platform [17]. Vascular cambium samples were collected under two conditions: from trees treated with a synthetic precursor of ethylene for 5 or 15 days and from control plants collected at the same intervals, as described by Junkes et al. [17].

In experiment (iii), RNA-Seq was carried out on two 15-month-old genotypes of Pra trees, resulting in 26 cDNA libraries. These samples were obtained from Bioforest SA in Concepción, Chile. Sequencing was performed using the Illumina GAIIx platform, as described by Carrasco et al. [15]. Stem tissue samples were subjected to inoculation (2, 6, and 12 days post inoculation (dpi)), wounding (2, 6, and 12 dpi), or no inoculation (0 dpi) for three different strains of *Fusarium circinatum* [15].

We established a unified bioinformatics pipeline tailored for analyzing these datasets, encompassing (i) quality filtering using Trimmomatic v.0.39 [37]; (ii) transcriptome assembly with Trinity v.2.11.0 [38]; (iii) transcript quantification with Salmon v.1.1.0 [39] using a k-mer size of 31; (iv) assembly evaluation with the Benchmarking Universal Single Copy Orthologs (BUSCO) v.5.1.2 tool [40]; and (v) transcriptome annotation with the Trinotate v.3.2.1 pipeline and the SwissProt database [41].

To determine the correspondence of the transcripts to the genomic regions containing probes, all the assembled transcripts were aligned to the LP scaffolds with the BLASTn v.2.11.0 tool [42]. The strict criteria included a minimum e-value of 1e-30 and a transcript coverage of at least 75%. To assign Gene Ontology (GO) terms and Enzyme Commission (EC) numbers to the assembled contigs, we considered the annotations obtained with the Trinotate pipeline.

### 2.5. Genome-Wide Association

To identify associations between genetic variants and SV-adjusted predicted least square means, we employed the Fixed and Random Model Circulating Probability Unification (FarmCPU) method, implemented in the rMVP R package v.1.0.6 [43]. In this analysis, we incorporated the first three principal components derived from a PCA conducted on the SNP dataset and the field sites where genotypes were assessed as fixed covariates. To ascertain SNPs significantly associated with phenotypic variation, we considered a threshold of 0.05, accounting for the Bonferroni correction. Furthermore, we expanded the marker set associated with SV based on estimates of LD, requiring a minimum Pearson correlation coefficient of 0.7.

For SNPs exhibiting significant associations with SV, functional inference was performed by examining neighboring genes and their putative annotations, which were obtained from the assembled transcriptome data. We established potential correspondences for each marker based on the alignments conducted with Pta, Pra, and Pel transcripts, considering a window of 10,000 bp on both sides of each SNP position, resulting in a total range of 20,001 bp.

### 2.6. SNP Prioritization through Machine Learning

In addition to the conducted GWAS, we explored the potential of the identified set of SNP markers in predicting the SV trait through a genomic prediction model. For this purpose, we employed a Bayesian ridge regression (BRR) approach implemented using the R package BGLR [44].

Considering our genotype dataset as a matrix *Z*_*n*×*m*,_ with *n* genotypes and *m* loci, the SNP effects (γ) were estimated considering the following equation:

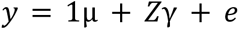

where *y* represents the SV trait, µ is the overall population mean, and *e is* the model residual 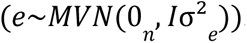. The SNP effects were assumed to follow a normal prior distribution 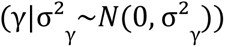 across all loci, considering the additive genetic variance σ^2^_γ_.

In contrast to predictions obtained using models trained with the entire set of markers, we explored the feasibility of feature selection (FS) techniques for subsetting SNP data based on putative phenotype‒genotype associations. We identified the intersection of at least two out of the three methods established [45–47]: (i) the gradient tree boosting (GTB) regressor model, (ii) Pearson correlation (maximum p value of 0.05), and (iii) L1-based FS with a linear support vector regression system (SVM). Using this subset, we assessed the importance of each SNP for prediction by calculating its feature importance with two different tree-based ML algorithms: a decision tree (DT) and a random forest (RF). We computed these estimates using the Gini index measure, which quantifies the reduction in the mean squared error associated with the prediction. Outliers of these measures were calculated by considering 1.5 times the interquartile range.

All the prediction models were evaluated considering a k-fold (k=10) cross-validation scenario, which was repeated 100 times and implemented with the scikit-learn Python v.3 module)[48]. The R Pearson correlation coefficient and mean squared error were calculated through the comparison of predicted and real values.

### 2.7. Gene Coexpression Networks

To evaluate the impact of phenotype-associated markers from a broader perspective, we constructed gene coexpression networks using Pel and Pra RNA-Seq data. We used the weighted gene coexpression network analysis (WGCNA) approach, implemented in the R package WGCNA [49]. After calculating a Pearson correlation matrix based on the expression of gene pairs, we estimated a soft power β for fitting each network into a scale-free topology. With the transformed correlations, we calculated a topological overlap matrix (TOM), which was converted into a dissimilarity matrix for the definition of groups of coexpressed genes using an average linkage hierarchical clustering approach together with adaptive branch pruning [49].

Following the identification of coexpressed groups containing genes surrounding SV-associated markers, we conducted an enrichment analysis for each gene set to identify enriched biological process GO terms. The topGO R package [50] was used, in combination with Fisher’s exact test and a multiple-comparison Bonferroni correction (p value of 0.01). Additionally, the REVIGO tool [51] was used to summarize and visualize the GO categories.

Based on the proportion of genes putatively associated with SVs within each coexpressed group, specific modules were selected for modeling an additional gene coexpression network using R statistical software [29]. The highest reciprocal rank (HRR) approach [52] was employed to investigate specific gene connections and network topology through centrality measures [53–55], including (i) betweenness, (ii) closeness, (iii) clustering coefficient, (iv) degree, (v) radiality, (vi) stress, and (vii) topological coefficient. The network was constructed considering Pearson correlations with a minimum coefficient of 0.8, and the 30 strongest connections for each gene were selected. Visualization of the coexpressed groups modeled by WGCNA in the HRR networks was performed using Cytoscape software v3.7.0 [56] and the R package igraph [57].

## 3. Results

### 3.1. Phenotyping and Genotyping

Seedlings derived from 308 LP families were planted across seven sites and analyzed in this study. Phenotypic analyses encompassed SV measurements obtained from 19,933 surviving individuals out of the initial 25,080 plants. The narrow-sense trait heritabilities varied between sites, ranging from 0.17 to 0.32, with an approximate mean of ∼0.22 observed at three of the seven sites (Table 1). Furthermore, the estimated dominance ratios ranged from 0.11 to 0.20.

**Table 1.** Estimates of individual narrow-sense heritability and dominance (± standard error) for stem volume trait at the sites evaluated.

| Site | Individuals | Families | Heritability | Dominance Ratio |
| --- | --- | --- | --- | --- |
| Site 1 | 3,588 | 307 | 0.22 (± 0.04) | 0.20 (± 0.06) |
| Site 2 | 3,600 | 307 | 0.22 (± 0.04) | 0.14 (± 0.05) |
| Site 3 | 3,545 | 307 | 0.18 (± 0.04) | 0.15 (± 0.05) |
| Site 4 | 3,588 | 308 | 0.17 (± 0.04) | 0.11 (± 0.05) |
| Site 5 | 3,588 | 307 | 0.32 (± 0.05) | 0.13 (± 0.06) |
| Site 6 | 3,600 | 307 | 0.18 (± 0.04) | 0.11 (± 0.07) |
| Site 7 | 3,571 | 307 | 0.22 (± 0.04) | 0.12 (± 0.05) |

Genotyping by sequence capture generated ∼996,611 million reads, which were subsequently aligned to the *P. taeda* reference genome. Given the significant occurrence of genomic duplications in *P. taeda*, we selectively focused on genomic regions associated with the 11,867 probes (± 500 bp) for SNP calling, merging regions located within 500 bp of each other. This process led to the identification of 10,412 sequences for alignment.

Approximately 84.5% of the sequenced reads were aligned to the designated probe sequences. This alignment process resulted in an initial count of 133,199 filtered SNPs, exclusively comprising biallelic SNPs, each with a minimum phred-scaled quality score of 20, located within sites having a minimum alignment score of 30, possessing a minimum allele frequency of 0.01, and no more than 30% of missing data. For the final SNP dataset employed in the GWAS, we applied more stringent criteria, allowing a maximum of 10% missing data and requiring both minimum and maximum allele frequencies of 0.05 and 0.95, respectively. These stringent filtering criteria resulted in a final set of 31,589 SNPs.

Regarding LD estimates between SNP markers, we observed rapid decay to half maximum (*R*^2^ = 0.05) occurring within less than 100 kb, as depicted in Fig. 1-i. A PCA conducted with the set of 31,589 SNPs revealed distinct genetic clusters corresponding to the crosses performed, as illustrated in Fig. 1-ii.

**Fig. 1.**
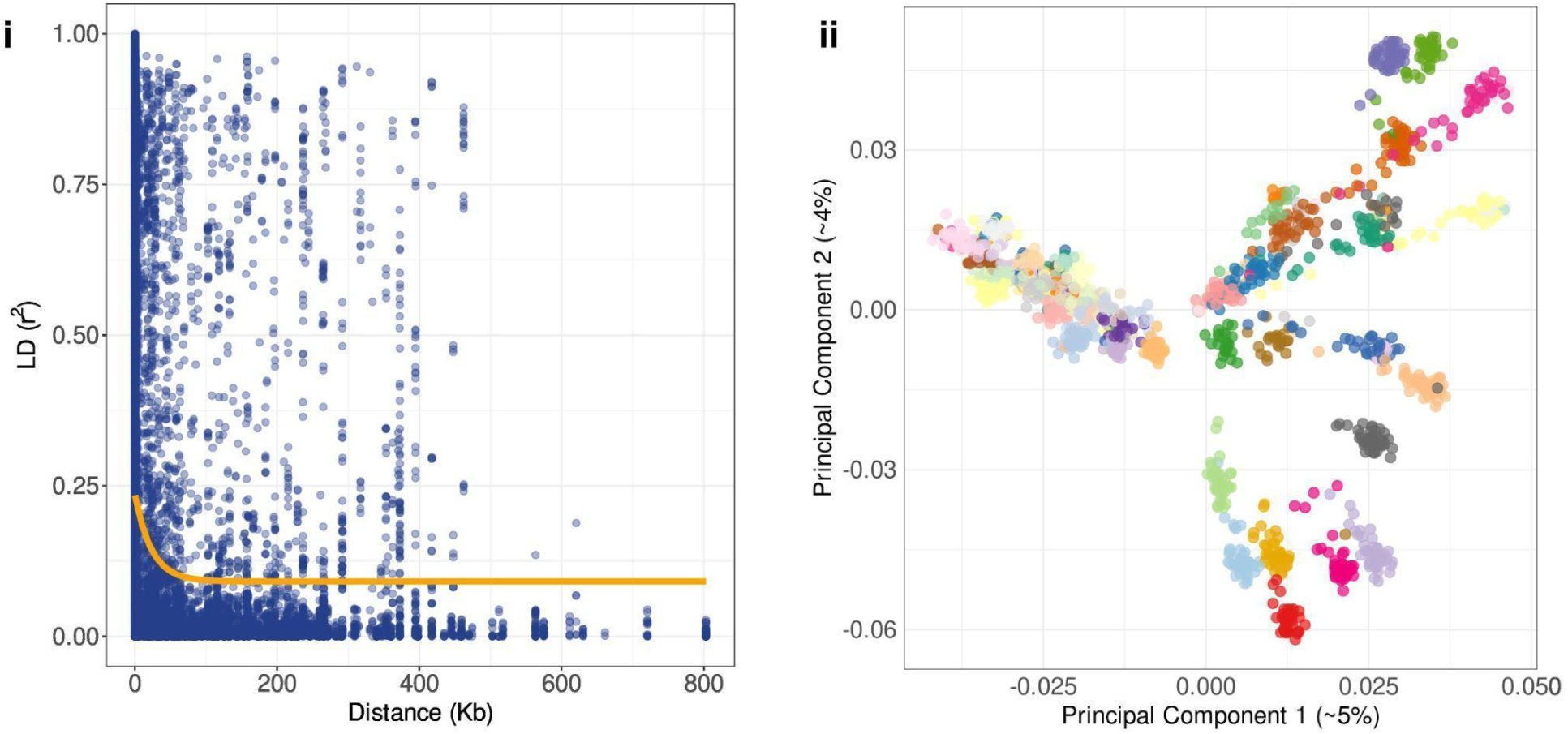
Genotypic analyses were performed on the SNP data and separated into (i) linkage disequilibrium (LD) decay considering squared Pearson correlations (R²) and distances calculated in kilobase pairs based on the respective probe sequences and (ii) principal component analysis (PCA) scatter plots of the *Pinus taeda* individuals, with colors representing their respective families (the first and second components explained approximately 5% and 4%, respectively, of the variance).

### 3.2. RNA-Seq Analysis

In our study, we utilized data from prior transcriptome experiments performed on Pta (60,899,426 reads), Pra (482,672,571 reads), and Pel (278,299,413 reads) to associate transcripts with the genomic regions containing probes. We obtained a significant number of RNA-Seq filtered reads for each species: 51,121,782 (∼83.94%) for Pta, 455,253,550 (∼94.32%) for Pra, and 276,114,142 (∼99.21%) for Pel. These reads enabled the assembly of 75,897 genes for Pta (comprising 146,652 isoforms with a median contig length of 772 bp and an N50 of 2,071), 137,103 for Pra (comprising 271,721 isoforms with a median contig length of 849 bp and an N50 of 1,921), and 50,781 for Pel (comprising 96,424 isoforms with a median contig length of 789 and an N50 of 1,499).

Notably, a substantial portion of the assembled contigs–78.1% for Pta, 98.4% for Pra, and 90.4% for Pel–could be recovered through BUSCO analysis, despite the presence of duplications attributed to the abundance of isoforms (62.1% for Pta, 97.4% for Pra, and 31.3% for Pel). Furthermore, approximately 40% of the total assembled genes were linked to SwissProt proteins via the Trinotate pipeline, with percentages of 43.77% for Pta, 41.64% for Pra, and 47.21% for Pel.

As our final set of assembled genes, we considered only those genes with expression in at least 75% of the samples within each dataset. This filtering process resulted in final gene counts of 38,472 for Pta, 37,499 for Pra, and 33,374 for Pel. Additionally, a notable proportion of the scaffolds containing probes–9,537 (92%) for Pta, 9,521 (91%) for Pra, and 8,132 (78%) for Pel–were associated with these genes.

### 2.3 GWAS

In our GWAS, we analyzed 1,692 LP genotypes characterized by 31,589 SNPs originating from 45 distinct families. The SV-adjusted predicted least square means were examined at four different sites. To account for the apparent population structure inherent to the families included in our study (Fig. 1-ii) and the influence of environmental effects, we integrated both sources of information as covariates in the GWAS model. Specifically, we incorporated the first three principal components obtained from the PCA performed on the SNP data and site information.

This comprehensive approach enabled the identification of a discernible pattern among the observed SNPs, revealing p values that deviated significantly from the null hypothesis, which assumes no association between SNP markers and SV variation (Fig. 2-i). Notably, seven SNP markers strongly correlated with SV variation were identified, each of which was located on distinct LP scaffolds (Table 2). Although a clear pattern of LD across these markers was challenging to ascertain due to the absence of a genomic reference at the chromosome level (Supplementary Fig. 1), we employed the Bonferroni correction as a stringent method to control for potential false positives arising from multiple comparisons.

**Fig. 2.**
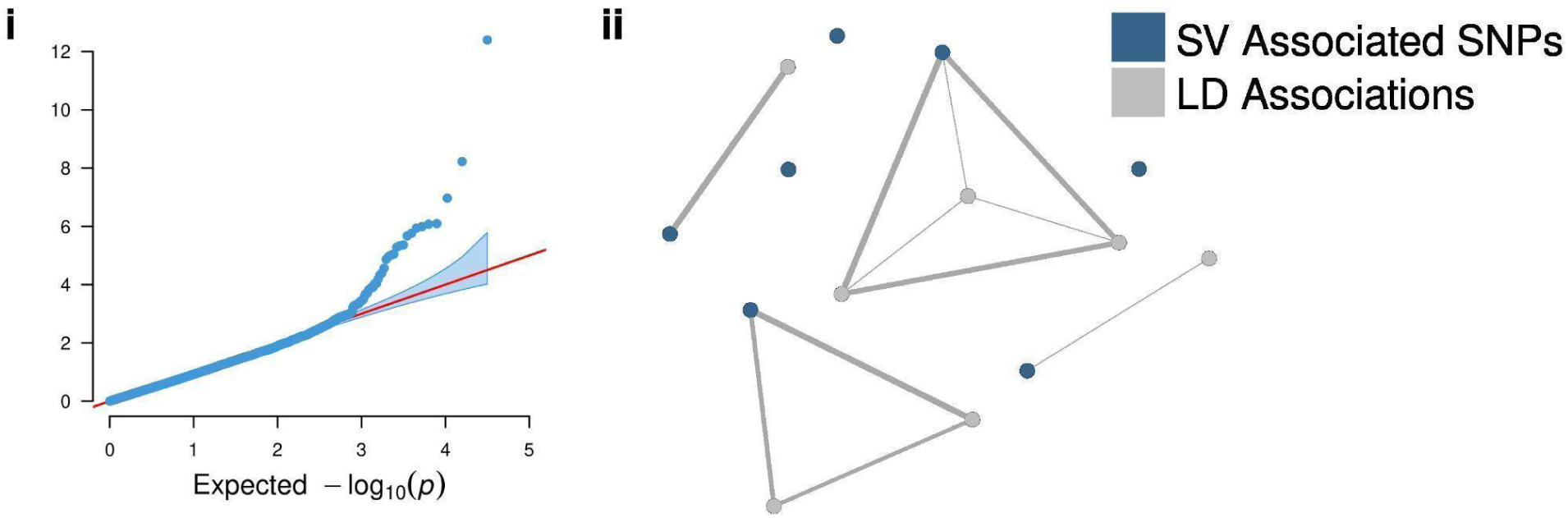
(i) Quantile‒quantile (QQ) plot of FarmCPU models in the GWAS with the stem volume (SV) trait. The set of markers associated with SV together with the other SNPs in linkage disequilibrium (LD) (minimum R Pearson correlation coefficient of 0.7) is displayed in (ii), which is a graph representing these associations within the set.

**Table 2.** A genome-wide association study (GWAS) revealed single nucleotide polymorphisms (SNPs). Gene annotations were included only if identified in at least two transcriptomes, considering *Pinus taeda*, *Pinus radiata*, and *Pinus elliottii*. The complete annotations are available in Supplementary Table 1. Underlined gene annotations are present in the *P. taeda* transcriptome, while those in parentheses are exclusively found in the *P. taeda* transcriptome.

| *Pinus taeda*<br>Sequence | Position | MAF | Effect | Transposons | Annotation of Surrounding Genes |
| --- | --- | --- | --- | --- | --- |
| super1496 | 457,445 | 0.11 | 0.0061 | 12 | Cysteine protease RD19A; Cysteine proteinase; and <u>Secreted RxLR effector protein 161</u> (<u>Shewanella-like protein phosphatase 2</u>) |
| super3461 | 42,253 | 0.09 | 0.0071 | 11 | – (<u>Secreted RxLR effector protein 161</u>) |
| scaffold11937 | 131,008 | 0.24 | -0.0033 | 6 | <u>Gag-Pol polyprotein</u>; and <u>Pro-Pol polyprotein</u> (<u>Auxin transporter-like protein 4</u>; <u>Auxin transporter-like protein 2</u>; and <u>Auxin transporter-like protein 1</u>) |
| scaffold67087 | 208,847 | 0.44 | 0.0030 | 11 | – (<u>Pro-Pol polyprotein</u>; and <u>Secreted RxLR effector protein 161</u>) |
| scaffold73604 | 618,978 | 0.06 | 0.0069 | 14 | <u>Secreted RxLR effector protein 161</u>; and <u>Zinc finger protein VAR3, chloroplastic</u> (<u>Putative ribonuclease H protein At1g65750</u> – Pta) |
| scaffold109055 | 54,249 | 0.05 | -0.0068 | - | – |
| scaffold187212 | 226,376 | 0.05 | 0.0065 | 6 | <u>Ubiquitin-60S ribosomal protein L40</u> |

In our investigation of the seven SNPs associated with SV, we identified five SNPs with positive effects on SV, contrasting with two that exhibited negative effects (Table 2). Notably, the minor allele frequencies remained below 15%, except for two markers, which exhibited frequencies of 24% (position 131,008 on scaffold11937) and 44% (position 208,847 on scaffold67087). These lower frequencies suggest the existence of specific allele configurations within certain families that may predispose individuals to SV variations.

Upon further investigation of the genes surrounding the SNPs identified through GWAS, we observed noteworthy similarities in the annotations derived from the transcriptomes of Pta, Pra, and Pel (Supplementary Table 1). The Pta transcriptome revealed 564 genes (1,218 isoforms) associated with GWAS-identified SNPs, while the Pel dataset revealed 65 genes (133 isoforms). In the case of the Pra transcriptome, 236 genes (830 isoforms) were found to be linked with the GWAS results. Except for a single genomic locus corresponding to SNP position 54,249 on scaffold109055, all the other regions demonstrated a substantial prevalence of transposable elements across the three utilized transcriptomes. This prevalence validates the extensive number of associated genes identified. The number of distinct transposable elements within each of these genomic regions ranged from 6 to 12, with a noteworthy abundance of retrovirus-related Pol polyproteins from transposon 17.6, retrovirus-related Pol polyproteins from transposon RE1, and retrovirus-related Pol polyproteins from transposons TNT 1-94 in these regions.

We uncovered additional annotations for genes neighboring the SNPs identified through GWAS. By specifically examining those associated with the Pta transcriptome or those simultaneously identified by Pra and Pel, we were able to identify significant proteins associated with these genes (Table 2). Notably, the secreted RxLR effector protein 161 was found to be linked with four out of the seven SNPs identified. Furthermore, we elucidated annotations for several other proteins, encompassing Shewanella-like protein phosphatase 2; cysteine protease RD19A; cysteine proteinase; gag-Pol polyprotein; pro-Pol polyprotein; auxin transporter-like proteins 1, 2, and 4; zinc finger protein VAR3, chloroplastic; putative ribonuclease H protein; and ubiquitin-60S ribosomal protein L40.

Due to the absence of a high-quality, chromosome-level genomic reference for LP, we incorporated an additional step into our analysis to identify SV-associated markers that were not detected in the GWAS results. Utilizing pairwise Pearson correlations between markers, we selected associations with a minimum correlation coefficient of 0.7 (Fig. 2-ii; Supplementary Fig. 2). This strategy allowed us to identify seven additional SNPs potentially associated with SV variations (Table 3; Supplementary Table 2). Subsequently, we linked 472 genes (comprising 1,004 isoforms) from the Pta transcriptome to these supplementary SNPs, with 459 genes also identified in the GWAS results. With respect to the Pel dataset, we discovered 48 genes (96 isoforms) linked to these markers, 46 of which were also identified via GWAS. In the Pra transcriptome, a total of 191 genes (with 725 isoforms) were associated with additional SNPs, and 181 of these genes were also identified in the GWAS.

**Table 3.** Linkage disequilibrium associations found with genome-wide association study (GWAS) results. Gene annotations were included only if identified in at least two transcriptomes, considering *Pinus taeda*, *Pinus radiata*, and *Pinus elliottii*. The complete annotations are available in Supplementary Table 2. Underlined gene annotations are present in the *P. taeda* transcriptome, while those in parentheses are exclusively found in the *P. taeda* transcriptome.

| *Pinus taeda*<br>Sequence | Position | MAF | Effect | Transposons | Surrounding Genes |
| --- | --- | --- | --- | --- | --- |
| super1496 | 457,299 | 0.11 | -0.0002 | 12 | Cysteine protease RD19A; Cysteine proteinase; and <u>Secreted RxLR effector protein 161</u> (Shewanella-like protein phosphatase 2) |
| super1496 | 457,451 | 0.11 | -0.0110 | 12 | Cysteine protease RD19A; Cysteine proteinase; and <u>Secreted RxLR effector protein 161</u> (Shewanella-like protein phosphatase 2) |
| super1496 | 457,550 | 0.07 | -0.0019 | 11 | Cysteine protease RD19A; Cysteine proteinase; and <u>Secreted RxLR effector protein 161</u> (Shewanella-like protein phosphatase 2) |
| scaffold49240 | 19,626 | 0.06 | -0.0009 | 1 | – |
| scaffold67087 | 208,716 | 0.44 | 0.0018 | 11 | – (Pro-Pol polyprotein; and <u>Secreted RxLR</u> <u>effector protein 161</u>) |
| scaffold67087 | 208,933 | 0.41 | -0.0013 | 11 | – (<u>Pro-Pol polyprotein</u>; and <u>Secreted RxLR effector protein 161</u>) |
| scaffold73604 | 618,878 | 0.06 | 0.0045 | 16 | <u>Secreted RxLR effector protein 161</u>; and <u>Zinc finger protein VAR3, chloroplastic (Putative ribonuclease H protein At1g65750)</u> |

Interestingly, one of the SNPs identified through LD analysis was found in association with a SNP from a different scaffold, diverging from the genomic location of the GWAS result (scaffold49240 at position 19,626 in the LD set associated with scaffold187212 at position 226,376 in the GWAS). This discrepancy is likely attributed to the utilization of a fragmented genomic reference and the substantial presence of transposable elements. Conversely, the additional SNPs retrieved through LD analysis demonstrated close proximity to the GWAS-associated SNPs. The distances, measured in base pairs, ranged from 6 (position 457,451 in super1496 associated with the GWAS result at position 457,445) to 146 (position 457,299 in super1496 associated with the GWAS result at position 457,445).

In terms of the effects of these LD markers, as estimated by the GWAS model, a consistent pattern emerged with the effects associated with the GWAS-identified SNPs. This consistency extended even to the LD association found on a different scaffold.

### 3.4. Identification of Trait–Marker Associations through Machine Learning

Due to the limited number of markers identified in association with SV variation, we employed an additional approach grounded in tree-based machine learning algorithms. Initially, we generated genomic prediction models using the BRR method, assessing the predictive capacity of the 31,589 SNPs in predicting the SV phenotype in a 10-fold cross-validation scenario. Our findings revealed a mean Pearson correlation coefficient (R) of 0.79 and a mean squared error of 0.00023 (Fig. 3-i), underscoring the robust predictive efficacy of the employed dataset for SV prediction.

**Fig. 3.**
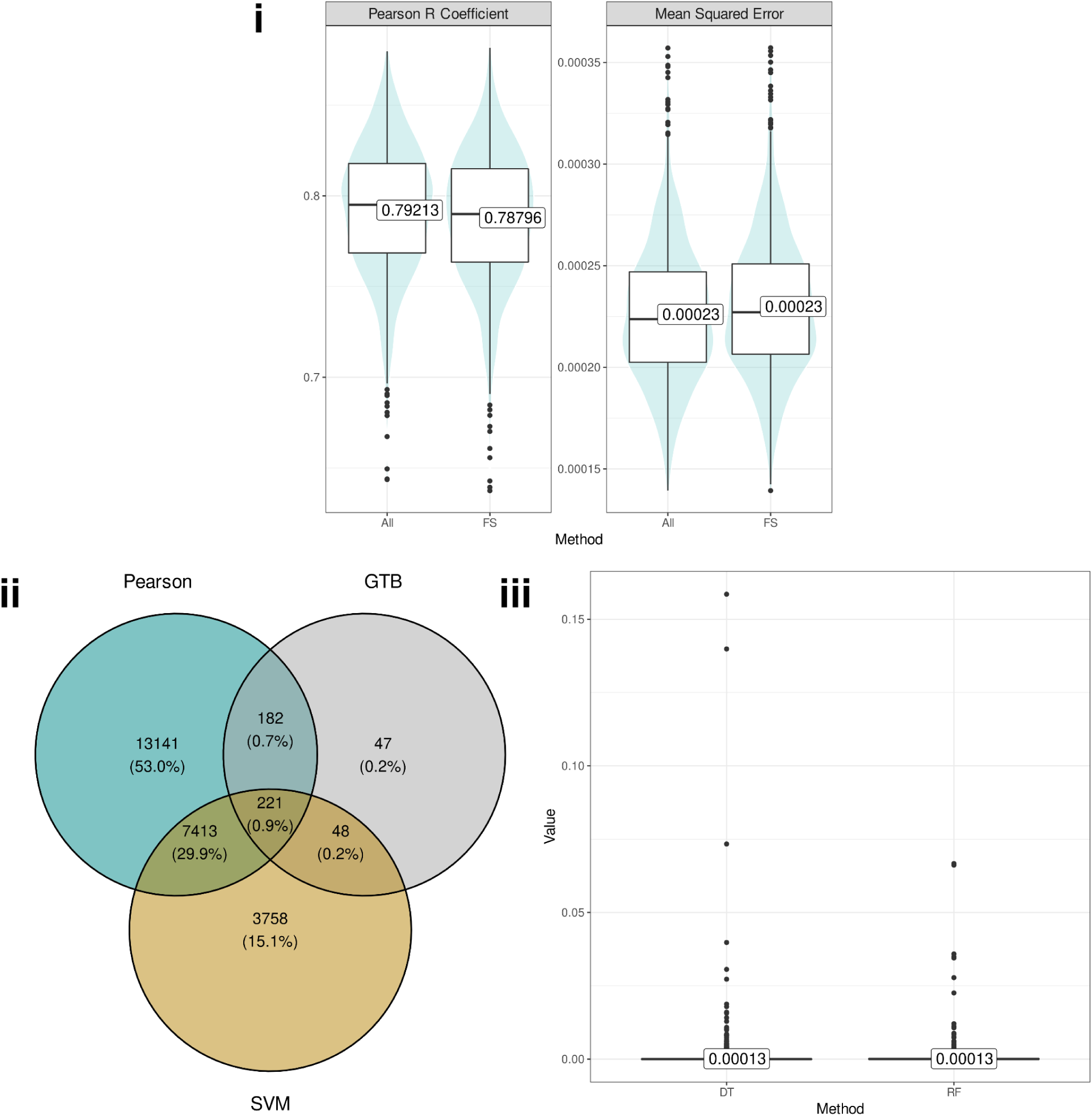
Genomic prediction results for stem volume using a total of 31,589 SNPs (All) and 7,864 selected through feature selection (FS) based on (a) Pearson correlations (p value of 0.05), (b) gradient tree boosting (GTB), and (c) support vector machine (SVM) analysis. (i) Predictive evaluations. (ii) Markers selected through each FS method. (iii) Distribution of the feature importance estimated for each of the 7,864 SNPs obtained through a decision tree (DT) and a random forest (RF) model.

In addition, different FS techniques were employed to reduce the dimensionality of the SNP data. Specifically, 498 SNPs were identified using the GTB algorithm, 20,957 SNPs through Pearson correlations, and 11,440 SNPs from the SVM model (Fig. 3-ii). To refine the selection process, we opted for a final subset of SNPs for estimating the SV predictive model. This subset was determined as the intersection of at least two out of the three FS methods, resulting in a set of 7,864 SNPs, representing approximately 24.89% of the initial marker data. Utilizing this reduced marker set, we achieved comparable accuracy in predicting SV (Fig. 3-i), indicating that a significant proportion of the phenotypic variance could be captured by this smaller dataset. The consistent predictive capacity observed between the total set of markers and the subset selected through FS, representing a reduction of approximately 75% in the initial SNP dataset, underscores the potential of SNP selection for genomic prediction. This process provides a streamlined set of markers for investigating phenotype‒genotype associations.

Finally, we utilized this selected dataset and employed tree-based algorithms to discern the relative importance of SNPs for predictive modeling, thereby elucidating additional phenotype‒genotype associations. The DT and RF models were employed, and Gini index estimates were obtained for each SNP (Fig. 3-iii). To focus on markers with a more substantial influence on prediction, we isolated outliers in these measures, identifying 1,506 SNPs for DTs and 635 SNPs for RFs. Our final selection of potential SV associations was established by determining the intersection between these two sets, yielding a refined set of 128 SNPs. Interestingly, it is noteworthy that all SNPs previously associated with GWAS were encompassed within this defined ML set.

Through the targeted selection of surrounding genes associated with the identified 128 SNPs, we identified an additional set of 2,333 genes in Pta, comprising 491 from GWAS, 414 from LD analysis, and 402 identified through both methods; these genes were distributed across 5,312 isoforms. In Pel, 327 genes were identified, with 50 from GWAS, 37 from LD analysis, and 36 from the overlap of both methods, spanning 738 isoforms. Similarly, in Pra, we found 1,113 genes, consisting of 222 from GWAS, 177 from LD analysis, and 173 from the convergence of both methods, distributed across 4,091 isoforms. Due to the substantial number of genes identified and the inherent limitations in statistical power associated with such extensive SNP selection, these genes were not individually examined. Instead, we chose to integrate these findings with the GWAS approach through gene coexpression networks, facilitating more comprehensive and holistic inference.

### 3.5. Gene Coexpression Networks

To conduct a comprehensive investigation of the molecular mechanisms underlying the genes associated with SV variation, we constructed gene coexpression networks using Pra (26 samples) and Pel (16 samples) data, which included 37,499 and 33,374 genes, respectively. Pta quantifications were excluded from the analysis due to the limited sample size (3 samples). Employing the WGCNA approach, distinct networks were modeled for each species, considering an estimated soft-power (β) of 14 for Pel (R² of ∼0.90 and mean connectivity of ∼53.10) and Pra (R² of ∼0.81 and mean connectivity of ∼143.20). In total, 251 groups were identified for Pra, ranging in size from a minimum of 81 to a maximum of 14,616 genes, with a mean of 471.79 genes per group. Similarly, 83 groups were established for Pel, varying in size from a minimum of 50 to a maximum of 4,858 genes, with an average of 132.96 genes per group.

Using the defined gene groups, we assessed the association of genes within each group with three distinct sets: (i) the 7 SNPs identified in the GWAS, (ii) the 7 LD associations with the SNPs from (i), and (iii) the ML-established set comprising 128 SNPs. Due to the more stringent criteria applied to the SNP sets in (i) and (ii), we specifically focused on groups featuring a minimum of two genes associated with these SNPs. Employing these criteria, we identified a total of 11 and 10 groups in the Pra and Pel networks, respectively. These groups exhibited enrichment in a diverse array of GO terms (Supplementary Table 3), encompassing processes related to gene expression regulation, defense response and signaling, metabolic processes, and plant development. This wide range of processes aligns with the known quantitative aspects of the SV trait.

In terms of the distribution of SV-associated genes within the selected modules, we observed a mean proportion of ∼1%, probably indicating specific genetic impacts on the biological processes performed by these groups. Notably, two groups in the Pra network (G39 and G68) exhibited a significant proportion of genes associated with the selected SV markers (∼92.78% of 194 genes in G39 and ∼57.69% of 78 genes in G68). These two groups, comprising 272 genes, were identified as a promising dataset for investigating the molecular mechanisms strongly associated with SV regulation; for this reason, these genes were used for the creation of a novel network via the HRR methodology.

This cohesive network comprised 270 genes (2 genes were discarded for not presenting significant correlations with the others) and a total of 3,008 connections. Although the network exhibited a distinct presence of two groups, potentially indicating two main functional profiles (Fig. 4-i), it was evident that these processes interacted through key gene associations. Analysis of a treemap generated from the biological process GO terms associated with this gene set (Fig. 4-ii) revealed involvement in processes such as DNA integration and recombination, cellular aromatic compound metabolic processes, cellular nitrogen compound metabolism, nitrogen compound metabolic processes, and transposition. Additionally, the network exhibited associations with viral latency and viral genome integration into host DNA and cells.

**Fig. 4.**
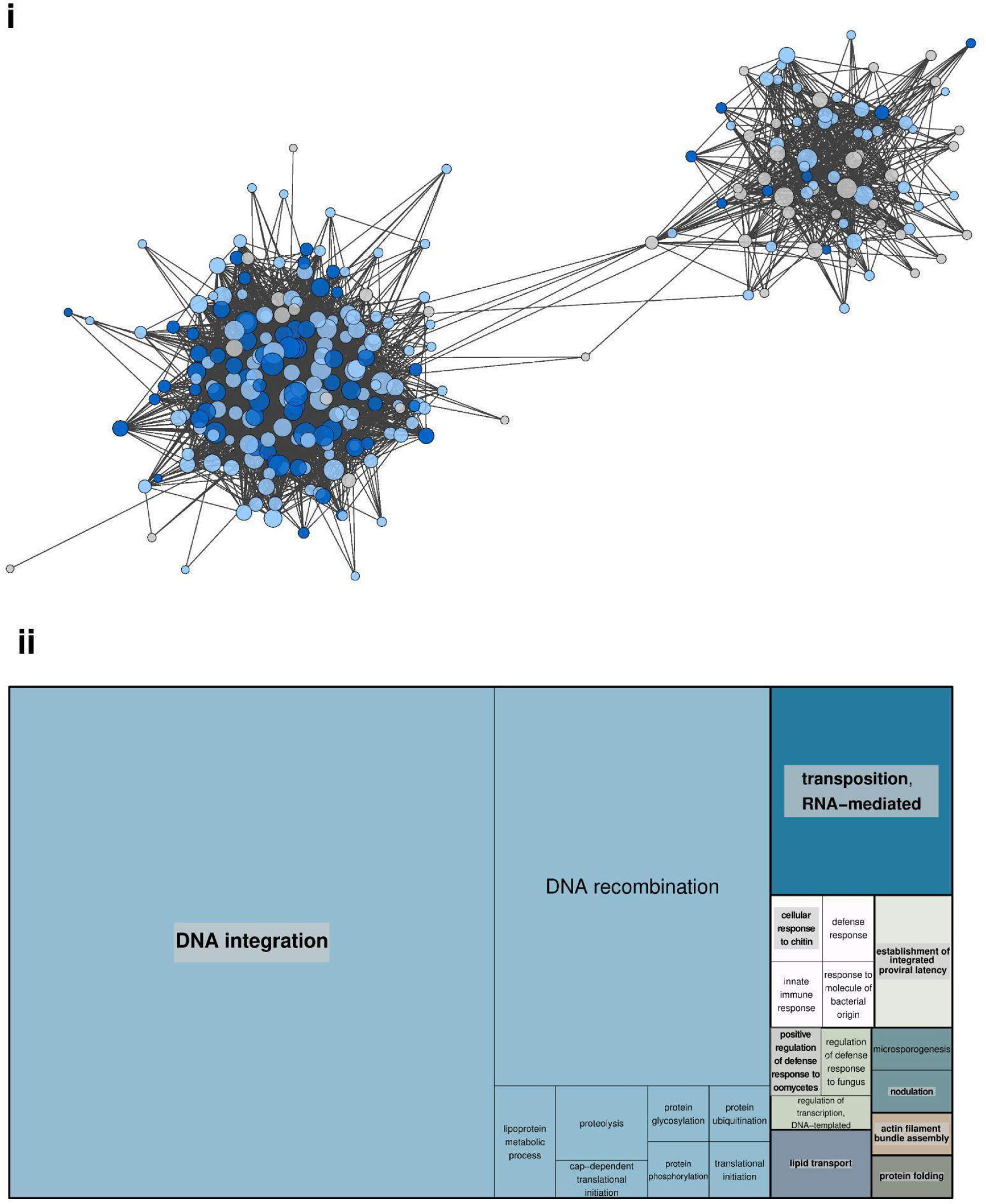
(i) Coexpression network created with the highest reciprocal rank (HRR) methodology considering groups of coexpressed genes selected in the weighted gene coexpression network analysis (WGCNA) modeled for *Pinus radiata* transcriptome data. Each node represents a gene, colored (a) gray (nondirect association with stem volume (SV) phenotype), (b) light blue (genes surrounding markers associated with SV through machine learning methodologies), and dark blue (genes of (b) and surrounding GWAS SNPs). (ii) Gene Ontology (GO) terms from the network summarized in a tree map.

In addition to identifying enriched GO terms, we explored the significance of key elements within the gene coexpression network through network centrality measures (Supplementary Table 4). A gene coexpression network serves as a mathematical representation of gene interactions, providing insights into the intricate relationships within a set of genes. Calculating gene centrality measures allows for the quantification of each gene’s importance in the network, reflecting its influence on the biological mechanisms executed by the network genes. Consequently, we focused on selecting genes with the top 5 values for the following centrality measures (Table 4): (i) betweenness, (ii) closeness, (iii) clustering coefficient, (iv) degree, (v) radiality, (vi) stress, and (vii) topological coefficient.

**Table 4.**
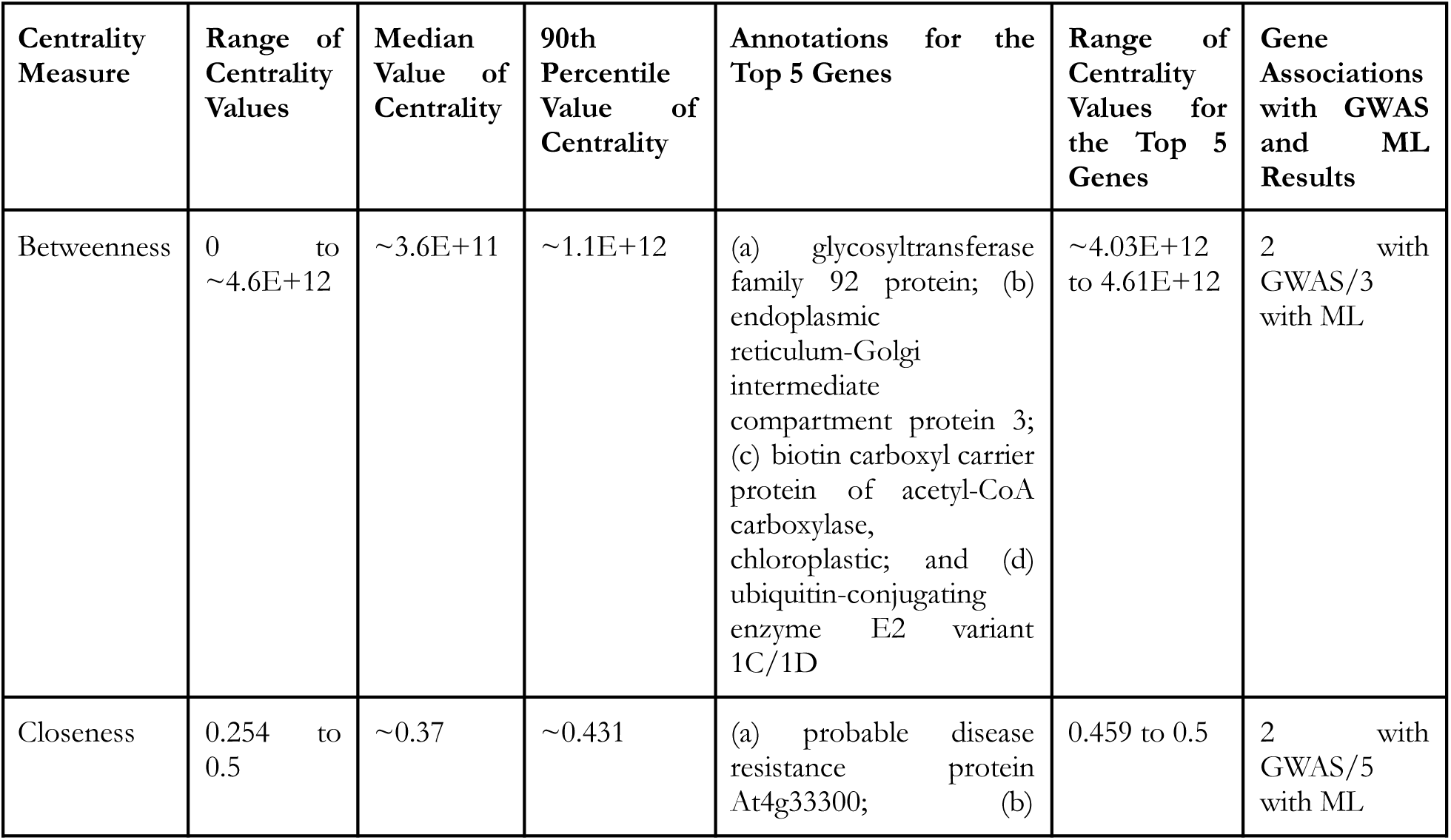
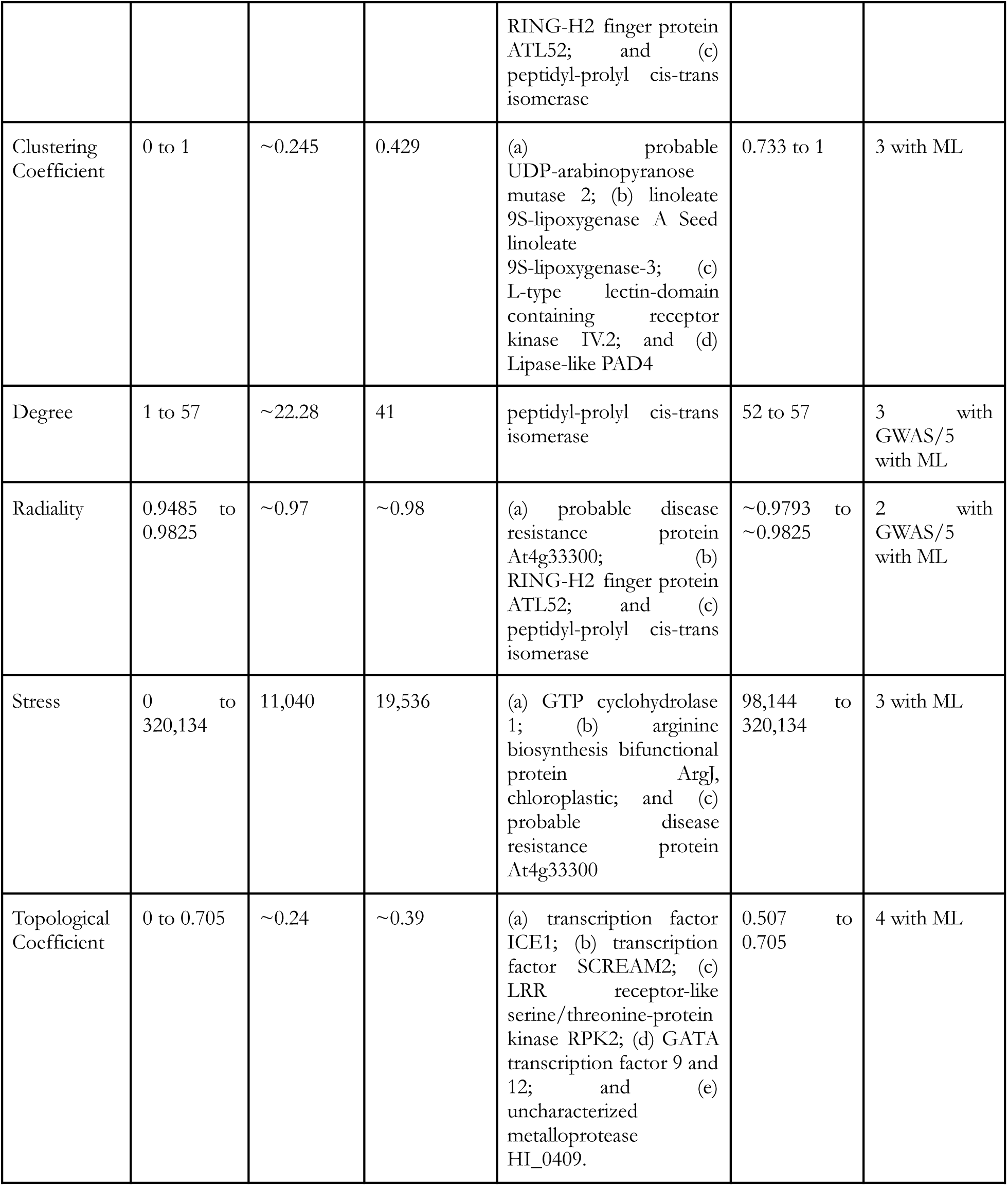
Identification of highly influential genes in the modeled gene coexpression network. The table presents the 5 genes with the highest values for each network centrality measure. Associated with potential annotations, these genes are also linked to results obtained from both genome-wide association analysis (GWAS) and machine learning (ML) approaches.

All of the genes within the subset of highly influential genes exhibited associations with the ML results. In contrast, for GWAS results, associations were not detected for only sets (iii), (vi), and (vii). This observation underscores the restrictive nature of GWAS analyses, characterized by a substantial number of false negatives due to p value correction. Notably, several gene annotations were consistently highlighted across centrality evaluations. For instance, the probable disease resistance protein At4g33300 was found in sets defined by measures of closeness, radiality, and stress; the RING-H2 finger protein ATL52 was found in sets defined by measures of closeness and radiality; and peptidyl-prolyl cis-trans isomerase was found in sets defined by measures of closeness, degree, and radiality.

Closeness centrality values serve as an indicator of how a gene can associate with others more quickly [54]. This finding aligns with the radiality measurement, which denotes the importance of a gene based on its centralizing role in the network [58]. Therefore, the concurrent association of these measures was expected. In the context of degree, the evaluation is based solely on the number of connections a gene possesses [53], which may or may not involve intermediate communication. When there is a high association with this intermediation aspect, stress measurements become a more suitable evaluation [54]. Despite their specific interpretations, there are similarities in how these measures calculate the influence of a gene within a network, leading to overlapping results.

In contrast, the betweenness, clustering coefficient, and topological coefficient evaluations revealed a more distinct profile among the ranked genes. The betweenness metric indicates a gene’s importance in facilitating its association [54]. Unlike closeness, neither clustering nor topological coefficients directly relate to the neighborhood of a node. While the clustering coefficient assesses a gene’s tendency to cluster with correlated genes, the topological coefficient reflects common neighbor associations [59]. This distinction sets them apart from the primary characteristics detected by the centrality measures of closeness, radiality, degree, and stress.

## 4. Discussion

Two key questions for forest tree breeders are which phenotypes should be measured and which trees should be selected [60]. Although the answer to this question depends on the location of the populations evaluated and the level of diversity involved, volume growth, a trait evaluated in this work, is one of the traits that forest tree breeders most often focus on for the evaluation and selection of phenotypes [10,20,61]. Since the 1960s and early 1970s, breeders have been able to produce 7–12% more volume per hectare at harvest than is possible with trees grown from wild seeds [62]. In our work, we performed an in-depth investigation of genomic associations with SV configurations, providing a rich characterization of not only such putative associations but also the genes and biological processes involved.

The selection of elite genotypes in LP populations can provide a gain in the volume of these trees of up to 50% over what is possible to produce with unimproved genotypes [63]. The response to selection is closely related to the heritability of the evaluated phenotype, and the greater this heritability is, the greater the response to selection [64,65]. However, in breeding programs that use genetic markers for selection, even to improve traits with low heritability values, such as volume, the efficiency of these markers can increase substantially. The narrow-sense heritability estimates for SV in LP vary considerably depending on the statistical strategies used and are substantially greater for more complex models [20,64]. We found heritability values for SV ranging from 0.17 to 0.32 among the seven sites analyzed, indicating that there are different trends of heritability at each site, but these trends are similar to those found by other authors [66,67].

While volume measurements constitute a prominent quantitative trait with a significant polygenic effect, identifying the genomic regions exhibiting the most pronounced effects on volume variation provides a starting point for unraveling the genetic architecture underlying tree growth. Although many GWAS results have not been directly translated into crop improvement strategies [68], it is well established that such outcomes lay the groundwork for subsequent genetic engineering approaches [9,69]. In contrast to the identification of introgressed alleles, the elucidation of favorable alleles associated with a specific trait facilitates more targeted selection of genes for genomics-based breeding [70]. Therefore, our study is important because it contributes to advancing the understanding of the molecular foundations that govern stem volume variation. This contribution, in turn, enables a comprehensive inference of the primary factors influencing this trait.

### 4.1. Genotype‒Phenotype Associations

Investigations employing GWAS in some pine populations have contributed valuable insights [71–74]. However, persistent challenges are evident, primarily related to the vast number of SNPs associated with complex quantitative traits and the limited comprehension of the underlying molecular mechanisms linking these genetic variants [75]. The identification of a gene influenced by a QTL is challenging, particularly in a species characterized by a genome as large as LP. Within our study, we identified genotypic markers associated with SV through GWAS, and noteworthy annotations were observed for genes and proteins that play pivotal roles in the development of LP volume.

In our exploration of the genetic basis of SV through GWAS, importantly, only allele differences exhibiting clear distinctions in stem volume were considered. Therefore, we aimed to identify genes intricately involved in the regulation of stem development. Notably, the outcomes of our GWAS revealed a significant association with cysteine proteinase. This protein plays a pivotal role in plant metabolism, influencing various aspects of plant physiology and development [76]. Consequently, we anticipate that genomic differences associated with the functions of cysteine proteinase may influence the phenotypic configuration of stem volume. Furthermore, Shewanella-like protein phosphatase 2 emerged as a direct association in our GWAS results. This protein is known for its role in transcriptional regulation, potentially impacting the expression of numerous other genes. Interestingly, a previous study revealed associations between the expression of Shewanella-like protein phosphatase 2 and genes encoding proteins involved in the photosynthetic machinery [77]. These noteworthy findings suggest a potential link between the genetic factors influencing stem volume and those involved in the intricate process of photosynthesis, shedding light on the coordinated regulation of plant growth and energy metabolism.

The ubiquitin-60S ribosomal protein L40 was also linked to our GWAS results. In addition to its involvement in the energy reserve metabolic process [78], this protein plays a crucial role in protein synthesis and the associated metabolic system. This multifaceted functionality enables plants to dynamically respond and adapt to various environmental challenges [79]. Understanding how plants respond to diverse stresses is pivotal for comprehending SV configurations, given the energy balance required for stress management and growth. Supporting the importance of the stress response in SV determination, our study also revealed the association of the cysteine protease RD19A with GWAS results. Previous research has linked this protease to water deprivation [80], a factor that directly impacts pine tree volume [81]. Furthermore, RD19A has been associated with responses to biotic stresses, including mechanisms affecting genes involved in resistance to *Ralstonia solanacearum* (RSS1-R) [82], a bacterium already described as responsible for extensive damage in forest trees [83].

Surprisingly, we identified a significant association between the GWAS results and the secreted RxLR effector protein 161, which is known for its role as a plant‒pathogen protein that manipulates plant defense responses by entering host cells and promoting virulence [84]. The association between LP genes and such proteins is attributed to the presence of the RxLR motif, which has already been suggested to be an indicator of plant immunity to these biotic stressors [85]. We believe that the evolutionary responses of plants to such stressors have shaped their genetic architecture, consequently leading to the development of the most favorable volume configurations in resistant genotypes. This intricate interplay between genetic factors and stress responses highlights the complexity of the regulatory mechanisms influencing stem volume in trees.

Although growth is recognized as a multifaceted trait influenced by several genetic factors [73,86], the GWAS conducted in our study revealed that only seven markers were significantly associated with growth. Santini et al. [73] argued that, despite employing high-throughput phenotyping techniques, growth-associated SNPs fail to account for a substantial portion of the observed phenotypic variation, which is an inherent GWAS limitation [87–89]. Consequently, we adopted ML-based models to complement our analysis and unveil additional genotype‒phenotype associations. Using the DT and RF models, we identified a more comprehensive set of 128 SNPs associated with the SV phenotype, providing valuable insights into the underlying genetic architecture. Notably, our findings revealed that the markers identified through GWAS were encompassed within this subset of 128 SNPs, reinforcing the robustness and efficacy of our hybrid methodology.

### 4.2. Insights into Molecular Mechanisms Governing Stem Volume Variation

Multiomics analyses have emerged as powerful tools for revealing the intricate genetic architecture underlying complex traits and mapping genetic variants with unknown biological roles to molecular mechanisms through integrative methodologies such as gene coexpression networks [26,86,90]. Considering that the phenotypic effect of each SNP selected by GWAS, LD, and ML can be very small, aggregating associated genes based on their expression patterns allows the identification of key physiological and regulatory processes within a set of genes [91]. Subsequently, loci identified as associated with a specific trait can be correlated with coexpressed modules within a network. Leveraging the ‘guilt-by-association’ principle [92], functional inferences can then be extended to the entire set of genes within these modules, providing a more comprehensive and robust understanding of functional relationships [26].

The genes and module networks associated with genotype‒phenotype relationships reveal a rich landscape of genetic interactions governing developmental processes in pine, consistent with findings from prior omics studies [93]. The intricate structure of these interactions poses a challenge for isolating individual genes for detailed characterization, given that changes in the expression of most genes are likely to exert some level of influence on multiple others. To address this complexity, we implemented a selection criterion focusing on associations within each module. This approach allowed us to narrow our focus to a reduced set of genes, emphasizing those with a more pronounced impact on SV alterations. By adopting this strategy, we aimed to increase the likelihood of uncovering genes that hold greater potential for providing in-depth insights and serving as promising targets for more extensive studies.

According to our investigation of the enrichment profiles of genes associated with biological processes within the selected gene groups, these genes likely play pivotal roles in the regulation of genetic material. Notably, transposable elements consistently occur throughout most network groups, highlighting their pervasive influence. Furthermore, the processes of DNA integration and recombination are strongly associated with alterations in DNA dynamics, particularly those favoring SV configurations. The identification of terms linked to defense responses strengthens the associations identified through GWAS, particularly those processes associated with the positive regulation of such responses. We infer that while the genetic architecture associated with SV encompasses a broad range of biological processes, the presence of a gene module strongly associated with SV alterations underscores a central regulatory set of genes influencing this diverse set of biological functions. This functional diversity corresponds to that of other network modules, emphasizing the interconnected nature of these regulatory processes.

Despite the shared functionality observed among the genes within the studied group, our investigation revealed distinct and specific roles for the genes in these subgroups. We identified highly influential genes based on ranked network centrality measures. Notably, peptidyl-prolyl cis-trans isomerase (PPIase) was the most highly connected gene in the network and exhibited direct associations with both the GWAS and ML results. This observation strongly suggested its regulatory role within the module. PPIases are known for their association with plant growth and development, and our findings reinforce their significance in this context. The genes associated with PPIases have been previously recognized as promising candidates for enhancing the abiotic stress tolerance of plants, a crucial aspect of both industrial processes and agricultural applications [94].

The measures of closeness, radiality, and stress identified additional genes with regulatory activity, albeit focusing on different aspects of network structure. Notably, our analysis underscores the importance of the RING-H2 finger protein annotation, a factor previously implicated in soybean seed weight and shape determination [95] and cold stress responses [96]. In alignment with the outcomes of our investigation, it is plausible that the genetic selection employed in pine tree breeding has promoted a mechanism involving DNA alterations in response to stress.

Furthermore, our study revealed annotations for GTP cyclohydrolase 1 and the arginine biosynthesis bifunctional protein ArgJ, neither of which were associated with GWAS or ML results. GTP cyclohydrolase 1 and ArgJ are integral to the biosynthesis of plant folate and arginine, respectively. Plant folate, an essential micronutrient, plays crucial roles in photorespiration and the synthesis of chlorophyll, plastoquinone, and tocopherol [97]. On the other hand, arginine, the primary source of nitrogen [98], is a pivotal factor influencing tree growth and development. Its acquisition, assimilation, storage, recycling, and efficient metabolic use significantly contribute to vascular development, forest tree productivity, and biomass production [99]. Even if not identified in association with SV genetic alterations, further exploration of these genes may offer valuable insights into targeted strategies for enhancing plant performance.

Furthermore, our investigation extended to genes contributing significantly to the overall gene interaction mechanism, as evidenced by the high betweenness centrality values of these genes. Notably, among these proteins is the biotin carboxyl carrier protein of acetyl-CoA carboxylase, a gene not highlighted by our GWAS or ML analyses. This protein is acknowledged for its mediation of stress responses, attributed to its involvement in fatty acid biosynthesis and lipid metabolism. Additionally, it has been associated with fundamental roles in plant growth and development [100]. Similarly, endoplasmic reticulum-Golgi intermediate compartment protein 3, which was identified by both GWAS and ML approaches, emerged as another prominent mediator. This protein actively participates in the trafficking and processing of proteins within the cell and is linked to driving defense responses [101]. The ubiquitin-conjugating enzyme E2, identified in association with both the GWAS and ML results, represents another key player in our network. This enzyme has been previously characterized for its pivotal role in protein degradation, exerting influence on cell cycle control and developmental processes [102]. Overall, we propose that these identified genes play crucial roles in mediating signal propagation for the configuration of volume, thereby influencing associated mechanisms through their mediating functions.

The genes demonstrating a local coordinated effect, as evidenced by their high clustering coefficient, are noteworthy candidates for coordinated regulation. Among them are the likely UDP-arabinopyranose mutase 2, which was identified in association with the ML results, and the lipase-like gene PAD4. UDP-arabinopyranose mutase 2 plays a vital role in plant growth and development [103], while the lipase-like gene PAD4 is important for its ability to impede pathogen growth and modulate defense responses [104]. These findings underscore the interconnected nature of the stress response and volume configuration, providing additional support for the association between these coordinated candidates and the underlying biological processes.

Ultimately, we employed the topological coefficient to evaluate the local interconnectedness of gene associations on a more extensive scale, complementing the insights provided by the clustering coefficient regarding local functional coherence. By selecting the ranked genes based on such a centrality measure, it is possible to provide insights into the mechanisms highly preserved in the network. Notably, our analysis highlighted the GATA transcription factor, which was also associated with the ML results. Despite not possessing the highest degree value in the network, the GATA transcription factor exerts a profound influence on the module. Its impact begins with a cohesive interaction among genes and extends to specific points within the broader network.

The GATA transcription factor, known to function as a growth suppressor based on previous research findings [105], exemplifies the potential of our investigation. The strength of its influence goes beyond local interactions, illustrating a cascade effect that reaches specific nodes in the network. This underscores the power of employing the topological coefficient to identify genes that, while not necessarily having the highest degree, significantly contribute to overall network dynamics.

Another transcription factor exhibiting a high topological coefficient in our coexpression network was ICE1, which was previously linked to the regulation of cold response and freezing tolerance [106]. Similarly, our analysis identified the transcription factor SCREAM2, which is known for its influence on the successive initiation, proliferation, and terminal differentiation of stomatal cell lineages [107]. Additionally, the LRR receptor-like serine/threonine-protein kinase RPK2 was found (which is also associated with ML results) to be clearly associated with stress response signaling [108]. All of these genes play pivotal roles in cellular regulation, emphasizing their importance within the coexpressed module.

Importantly, the regulatory functions attributed to these genes, as evidenced by their high topological coefficients, warrant comprehensive evaluation together with other analytical outcomes. Employing an integrated approach ensures a holistic comprehension of the regulatory landscape within the coexpressed module, unveiling potential interactions and coordination among these genes. Our findings highlight the limitations inherent in relying exclusively on GWAS for the identification of key elements within gene modules, reinforcing the efficacy of machine learning (ML) techniques in uncovering relevant genes that may elude traditional GWAS approaches. Despite the absence of direct GWAS associations, our integrative approach, in which multiple biocomputational methods were combined, enabled a comprehensive exploration of the regulatory mechanisms associated with SV variation. Our study not only offers insights into the SNP markers identified but also delineates a subset of genes characterized by unique regulatory features. This subset represents a compelling avenue for in-depth investigations, underscoring its significance for future research endeavors.

## Acknowledgments

We acknowledge Matias Kirst from the Forest Genomics Laboratory at the University of Florida for providing us with the data analyzed in this research.

