## Supplementary Figures for "Genome-wide association reveals novel insights into the molecular mechanisms regulating stem volume in *Pinus taeda*"

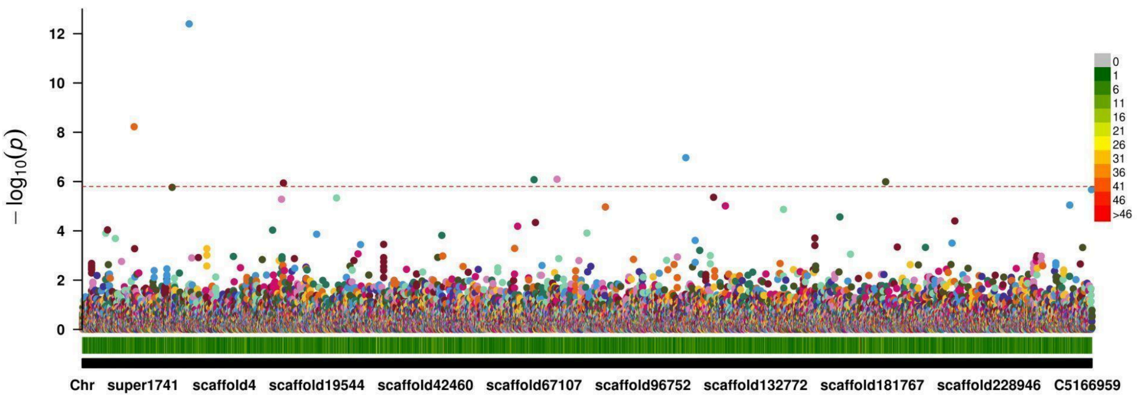


**Supplementary Fig. 1.** Manhattan plot for the p values derived from the genome-wide association study (GWAS) conducted to identify SNP associations with stem volume variation in *Pinus taeda*. Each data point in the scatter plot represents an SNP, with color coding based on the respective scaffold location.

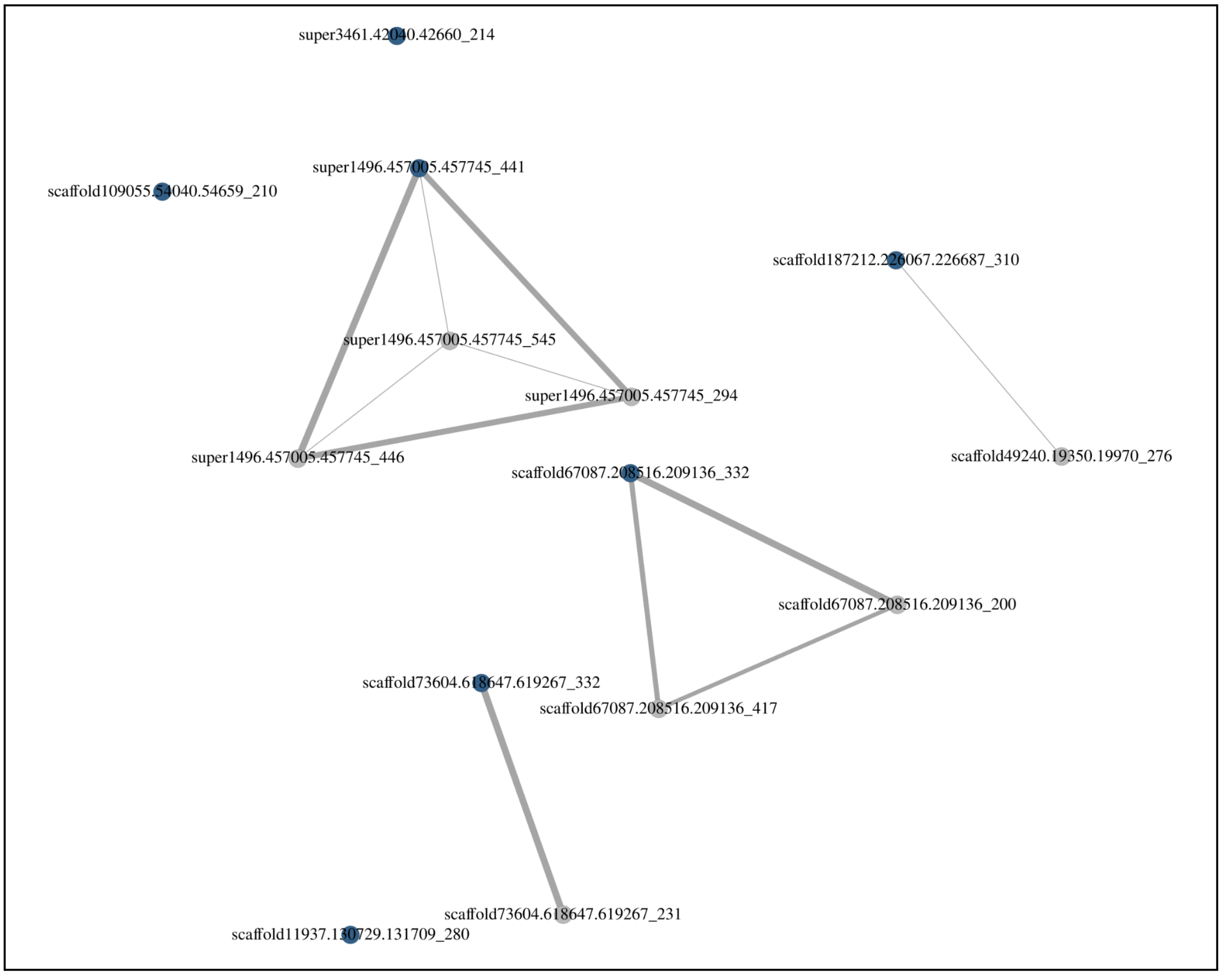


**Supplementary Fig. 2.** Linkage disequilibrium (LD) associations of single nucleotide polymorphisms (SNPs) in LD (minimum R Pearson correlation coefficient of 0.7) with markers identified through genome-wide association study (GWAS) analysis. Each point in the graph corresponds to a SNP, distinguished by color: blue indicates SNPs associated with stem volume identified through GWAS, while gray represents SNPs associated through LD.
